# Enhancing Secondary School Students’ Attitudes towards Biology through Guided Inquiry-Based Lab Instructions

**DOI:** 10.1101/2025.05.29.656923

**Authors:** Ashebir Mekonnen Chengere, Beyene Bobo Dobo, Samuel Assefa Zinabu, Kedir Woliy Jilo

## Abstract

Attitude plays a vital role in achieving modern science education goals by enhancing students’ conceptual understanding, motivation, academic performance, interest, and engagement in scientific inquiry. However, in many Ethiopian schools, attitudes toward science are often overlooked, and traditional teacher-centered instruction combined with limited resources hampers their development. To address these issues this study examined the effect of guided inquiry-based laboratory experiments enriched instruction (GIBLEI) on secondary school students’ attitudes toward biology, using a quasi-experimental design with pretest-treatment-posttest phases. Two classes from selected schools were randomly assigned to an experimental group (EG, N=46) and a control group (CG, N=29). Over eight weeks, the EG received GIBLEI, while the CG experienced traditional laboratory experiments enriched instruction (TLEI). Attitudes were measured using a 5-point Likert scale. The results showed that GIBLEI significantly improved students’ overall attitude towards biology, enthusiasm for biology, perception of biology as a course, and understanding of biology as a process, compared to TLEI. However, it did not affect their views of biology as a career. GIBLEI also promoted gender inclusivity by reducing attitude differences between male and female students. These findings highlight the benefits of GIBLEI in fostering positive attitudes, engagement, and inclusivity in biology education, enhancing both student outcomes and equity in science learning.

## 1. Introduction

The development of scientifically literate individuals is indispensable for the advancement of societies, impacting not only their socio-economic fabric but also their scientific progress (1).Consequently, significant attention has been directed towards enhancing science education, particularly at the secondary level, across various nations. Notably, countries in Sub-Saharan Africa, including Ethiopia, have embarked on curriculum reforms, transitioning from knowledge-centric to competence-based approaches. This shift aims to equip learners with the essential attitudes, skills, and values in science teaching, thereby fostering a trajectory towards a sustainable future(2).

In this context, attitude becomes a central construct in efforts to evaluate and improve biology education. It refers to students’ opinions, emotional dispositions, and cognitive frameworks that influence how they perceive and respond to biology as a subject (3). Educational researchers have conceptualized attitude in multiple dimensions, including students’ interest in the subject, perceived relevance, aspirations for biology-related careers, and views on instructional methods (4;5).These dimensions provide a structured lens for analyzing both positive engagement and negative perceptions that may hinder achievement and participation, in settings like Ethiopia, where such challenges remain underexplored.

Laboratory experiments play a crucial role in enhancing student engagement and improving attitudes toward biology by offering hands-on, experiential learning that connects abstract concepts to real-world experiences (6). However, many biology classrooms still rely on teacher-centered, traditional laboratory methods, where students follow rigid, step-by-step instructions with predetermined outcomes, limiting creativity and opportunities for independent thinking. To improve students’ attitudes toward biology, it is essential to transition from these traditional methods to more student-centered, inquiry-driven approaches that foster autonomy and engagement. Additionally, addressing gender biases and promoting inclusivity in science education can further enhance students’ attitudes by creating a more equitable learning environment, where all students, especially girls, feel empowered to pursue their interests in biology and other STEM fields (7).

Inquiry-Based Learning (IBL) has emerged as a transformative pedagogical strategy that not only enhances students’ conceptual understanding but also significantly improves their attitudes toward science subjects, including biology. By placing students at the center of the learning process, IBL encourages them to ask questions, explore real-world problems, construct their own knowledge, and reflect on their learning (8). This active engagement fosters a sense of curiosity and ownership, which contributes to more positive attitudes towards the subject matter. Several studies have demonstrated that IBL can increase students’ interest, enjoyment, and motivation in learning biology by making the subject more relevant, interactive, and intellectually stimulating (9). Moreover, hands-on, inquiry-based laboratory activities allow learners to connect theoretical concepts to practical experiences, which has been shown to reduce negative perceptions and anxiety associated with science learning (10). In biology education specifically, IBL has been associated with increased enthusiasm and engagement, which are key dimensions of student attitude (4). Empirical evidence further supports these outcomes; for instance, Minner et al (11) found that students in inquiry-based environments not only developed stronger understanding but also expressed more favorable attitudes toward science compared to those taught through traditional, didactic instruction.

Implementing Inquiry-Based Learning (IBL) in resource-constrained contexts like Ethiopia presents several challenges that hinder its effectiveness in fostering positive student attitudes toward biology. Key barriers include inadequate laboratory facilities, limited access to instructional materials, and a shortage of teachers trained in inquiry methodologies (12). These constraints make it difficult to conduct hands-on, exploratory activities central to IBL, often causing students to view biology as abstract and disconnected from real life. Additionally, the open-ended nature of IBL may overwhelm students who lack foundational knowledge or prior exposure to student-centered pedagogies, potentially leading to confusion, frustration, and decreased interest (13). Teachers, many of whom are accustomed to traditional, lecture-based methods, may resist implementing IBL due to limited professional development and discomfort with facilitating student-directed learning. In environments where students are unaccustomed to taking initiative, the absence of scaffolding can further weaken engagement and hinder the development of favorable attitudes toward biology(8). Similar challenges have been documented in other developing nations such as Kenya, Nigeria, Ghana, and Bangladesh, where IBL is undermined by large class sizes, inadequate resources, and insufficient teacher preparation(14;15;16;17).

To address these limitations, this study introduces the Guided Inquiry-Based Laboratory Experiments Enriched Instructional (GIBLEI) approach, which adapts inquiry pedagogy to low-resource settings by providing structured, teacher-supported investigations. Unlike fully open-ended IBL, GIBLEI follows the Guided Inquiry-Based Laboratory Instruction (GIBLI) model, offering students a defined research question and step-by-step guidance through experimentation, analysis, and reflection (18). This approach combines cognitive apprenticeship, where teachers model scientific thinking and gradually transfer responsibility, with structured scaffolding, supporting learners’ autonomy, competence, and relatedness as emphasized in Self-Determination Theory (19). GIBLEI also enhances content relevance by linking biology to real-world contexts, which helps sustain interest and emotional engagement. It encourages collaboration, problem-solving, and peer interaction, consistent with constructivist learning principles that promote inclusive participation, particularly for marginalized groups such as female students (20). By integrating virtual labs, using locally available materials, and incorporating professional development for teachers, GIBLEI offers a feasible and impactful alternative to traditional methods, improving both conceptual understanding and students’ attitudes toward biology (21).

This study empirically investigates the effectiveness of the Guided Inquiry-Based Laboratory Experiments Enriched Instructional (GIBLEI) approach compared to the Traditional Laboratory Experiments Enriched Instruction (TLEI) method in shaping secondary school students’ attitudes toward biology in North Shoa Zone, Oromia Regional State, Ethiopia. Specifically, it examines students’ attitudes across four key dimensions: enthusiasm for biology, perceptions of biology learning, understanding of biology as a process, and interest in biology as a career. These dimensions represent a multidimensional framework that enables both overall and domain-specific analyses of instructional impact. To assess the effects of the instructional methods and identify any significant differences between the experimental (GIBLEI) and control (TLEI) groups, a set of null hypotheses was formulated.

**Ho**_**1**_: There is no significant difference between the overall attitude mean scores of students taught using the GIBLEI approach and those taught using the TLEI approach.

**Ho**_**2**_: There are no significant differences between the mean scores of students on the four dimensions of attitude (enthusiasm toward biology, biology learning, biology as a process, and biology as a career) between the two instructional groups.

**Ho**_**3**_: There are no significant differences between pre-test and post-test attitude scores within each instructional group.

**Ho**_**4**_: There are no significant differences in overall attitude or attitude dimensions between male and female students within each group.

## 2. Materials and Methods

### 2.1 Research Design

This study examined the impact of the Guided Inquiry-Based Laboratory Experiments Enriched Instructional (GIBLEI) approach on grade 10 students’ attitudes toward biology. A total of 75 students (35 boys and 40 girls) from two public secondary schools in Fitche Town, North Shoa Zone, Ethiopia, participated in the study. The experimental group (n = 46), drawn from Fitche Secondary School, received instruction through the GIBLEI approach, while the control group (n = 29), from Abdisa Aga Secondary School, was taught using the Traditional Laboratory Experiments Enriched Instructional (TLEI) method (see Table 1). The intervention targeted Unit 4 of the grade 10 biology curriculums, Food Making and Growth in Plants, which includes plant organs, photosynthesis, transport, and response mechanisms, topics often associated with persistent student misconceptions (22).

**Table 1.** Layout of the research design.

| Group |  | Intervention Sequence |  |
| --- | --- | --- | --- |
| Experimental | Pretest | X<sub>1</sub> (GIBLEI Approach) | Post-test |
| Control | Pretest | X<sub>2</sub> (TLEI Approach) | Post-test |

Note: X<sub>1</sub> =Guided inquiry based laboratory experiments enriched instructional (GIBLEI) approach
X<sub>2</sub> =Traditional laboratory experiments enriched instructional (TLEI) approach

A non-equivalent quasi-experimental pretest-posttest design was employed. This design is appropriate for educational contexts where random assignment is impractical due to the use of intact classrooms (23). To enhance group comparability, schools were selected based on the similarity of their facilities and teacher qualifications. Pretests were administered to both groups to control for baseline differences and improve the reliability of post-intervention comparisons.

### 2.2 Samples and Sampling Technique

This study employed a multistage sampling technique to select participants. In the first stage, two public secondary schools, Fitche Secondary School and Abdisa Aga Secondary School, were purposively selected from the North Shoa Zone of the Oromia Region, Ethiopia. Efforts were made to ensure that the schools were comparable in terms of infrastructure, teaching staff, and resources, as differences between groups can be a concern in non-equivalent quasi-experimental designs (24) Both schools are located in the urban area of Fitche Town and have similar facilities, including laboratory chemicals, equipment, library resources, and ICT infrastructure. In the second stage, the schools were randomly assigned to either the EG (Fitche Secondary School) or the CG (Abdisa Aga Secondary School). In the third stage, two highly qualified and experienced biology teachers were purposively selected. Both teachers hold a Master’s of Science degree in Biology and have over 14 years of teaching experience. Finally, a grade 10 class was randomly selected from each school and assigned to the EG or CG, ensuring fairness in group allocation. The EG consisted of 46 students (22 male, 24 female), and the CG included 29 students (13 male, 16 female), totaling 75 grade 10 students (35 boys and 40 girls).

Moreover, the study focused on Unit 4 of the Grade 10 biology curriculum, “Food Making and Growth in Plants,” which covers topics such as plant organs, leaves, photosynthesis, transport, and response mechanisms. This unit was purposefully selected because it addresses common student misconceptions, such as the misunderstanding that plants require only carbon dioxide, and confusion about the processes of photosynthesis and respiration (22).

### 2.3 Data collection instrument

The Biology Attitude Questionnaire (BAQ) was adapted from Kaur & Zhao (5) assess secondary school students’ attitudes toward biology. The questionnaire, consisting of 36 items, was administered both before and after an intervention to capture changes in students’ perceptions. A five-point Likert scale measured attitudes across four dimensions: enthusiasm toward biology, biology learning, biology as a process, and biology as a career. Responses ranged from “strongly disagree” (1) to “strongly agree” (5), with positive items scored directly and negative items reverse-scored to maintain consistency. Mean scores for each dimension were calculated by averaging responses to the related statements, allowing for detailed insights into students’ attitudes.

The first dimension, enthusiasm toward biology, includes seven statements reflecting students’ feelings toward the subject, ranging from enjoyment and importance (positive extreme) to aversion (negative extreme). The second dimension, biology learning, encompasses 11 statements that explore students’ perspectives on discovery, comprehension, and active engagement (positive views) versus formal, limited, and mechanical learning (negative views). The third dimension, biology as a process, contains nine statements addressing the students’ understanding of biology as a dynamic, evolving discipline (positive view) versus a rigid, nearly complete field (negative view). Lastly, the fourth dimension, biology as a career, includes nine statements assessing perceptions of biologists and interest in biology-related careers, contrasting positive views of biologists as competent professionals with negative stereotypes of them as antisocial or eccentric. By averaging mean scores across all 36 items, the overall attitude toward biology was determined, while individual dimension scores provided a more granular understanding of students’ perspectives. This structured approach facilitated a comprehensive analysis of changes in attitudes, offering insights into how students perceive biology and its relevance to their lives and future aspirations.

#### 2.3.1 Validity of the BAQ

Validity refers to the extent to which an instrument measures what it is intended to measure (25). In this study, particular emphasis was placed on content and face validity to ensure that the Biology Attitude Questionnaire (BAQ) accurately captures students’ attitudes toward biology. The BAQ was adapted from the Physics Attitude Questionnaire developed by Kaur & Zhao (5), with subject-specific terminology changed from physics to biology and irrelevant items removed. Content validity was established through a systematic review by a panel comprising PhD students specializing in biology education and experienced biology teachers. The original scale included 60 items across five dimensions: Enthusiasm toward Physics, Physics Learning, Physics as a Process, Physics Teacher, and Physics as a Future Vocation. A collaborative refinement process, involving the biology teachers and PhD students, streamlined the scale to 36 items grouped into four dimensions: Enthusiasm toward Biology, Biology Learning as a Course, Biology as a Process, and Biology as a Career. These experts evaluated the revised items to ensure clear alignment with the four key attitude dimensions: Enthusiasm toward Biology, Biology Learning as a Course, Biology as a Process, and Biology as a Career.

Face validity was ensured through a structured evaluation by the same group of experts. Printed copies of the BAQ were distributed, and the experts assessed each item for clarity, simplicity, and cultural relevance to secondary school students. They also evaluated whether the items appeared to effectively measure attitudes toward biology and provided feedback on any ambiguities or potential misunderstandings. Group discussions were then conducted to address these issues and refine the items collaboratively. This process enhanced the perceived clarity and relevance of the BAQ, ensuring it was appropriate and understandable for the target audience.

To make the BAQ accessible to all participants, it was translated into Amharic and Afan Oromo by language experts. To validate these translations, a back-translation process was conducted in which the items were retranslated into the original language by independent translators. These back-translated versions were compared with the original questionnaire to identify and resolve any discrepancies in meaning. Experts in psychology and language studies reviewed the translated versions to ensure accuracy, clarity, and cultural relevance, confirming that the instrument maintained its validity across different languages and was suitable for diverse linguistic backgrounds within the Ethiopian context.

#### 2.3.2 Reliability of the BAQ

Internal consistency reliability, commonly measured by Cronbach’s coefficient alpha, assesses how well a set of items measures a single construct (26). In this study, Cronbach’s alpha was used to evaluate the internal consistency of the Biology Attitude Questionnaire (BAQ). The pilot study yielded an overall alpha coefficient of 0.834, indicating good reliability. However, four items (2, 16, 22, and 28) were removed as they negatively affected the internal consistency of the questionnaire. After these adjustments, the revised BAQ demonstrated reliable internal consistency across its four attitude dimensions: “Enthusiasm” (α = 0.701), “Biology Learning” (α = 0.703), “Biology as a Process” (α = 0.707), and “Biology as a Career” (α = 0.704). These alpha values fall within the acceptable range, suggesting the questionnaire is reliable for capturing attitudes toward biology. The four removed items were carefully reviewed to address their initial issues with internal consistency and ensure they met reliability standards. After this review and refinement, these items were integrated into the improved version of the Biology Attitude Questionnaire (BAQ). The finalized version of the BAQ, incorporating these adjustments, was validated and prepared for the actual data collection phase, ensuring it served as a reliable and robust tool for assessing student attitudes toward biology.

### 2.4 The treatment procedures

#### 2.4.1 Pre-Treatment Procedure

On the 1^st^ stage, the teacher and the laboratory technician from the EG were given a training on the instructional and training materials prepared by the researchers. The training included a detailed description of the GIBLEI. It also encompassed how to use the 5E lesson plan format to prepare a daily lesson plan by teachers. The training was given by the researchers and lasted for 3 days, 90 minute per day. The first day, 90 minutes were used for explanation on the materials and the rest days were for practice in actual classroom. In addition, orientation was given on the aim of the research to the teacher and laboratory technician from the CG before intervention. Then, the BAQ pretest was administered to the two groups. After the completion of all the preliminary activities, intervention was started. The intervention was conducted over approximately eight weeks, spanning from 25/03/2023 to 30/05/2023 during the second semester of the 2023 Ethiopian academic year. In the syllabus it was shown that the unit needed 24 periods to be accomplished. The EG was taught using GIBLEI approach, whereas the TLEI approach was employed for the CG. With the support of the researchers and laboratory technicians, the selected biology teachers delivered the following sub-topics from Unit 4 of the Grade 10 Biology textbook to both the EG and the CG. These were the organs of a flowering plant and the leaf, photosynthesis, transport in plants, and response in plants.

#### 2.4.2 Treatment procedure

The intervention began after all preparatory tasks were finished. The procedures listed below were applied to every GIBLEI. The procedure was modified based on Blanchard et al (27). With the lab technician’s assistance, the chosen teacher guides every step of the process. The following procedures were followed for all the 19 practical activities.

1. A week prior to each class, students were given semi-structured problems from unit 4 of the biology curriculum for grade 10.
2. The students used to look for an experimental procedure and plan their experiments for every problem until the following lab practice.
3. They used the results of their search to determine the layout of their experiment.
4. The students used to share their ideas and discuss their experimental design with other groups. At this point, they would go over every step of the experiment, the materials, and the rationale behind the materials and procedure they had selected.
5. The lab technician and the teacher have supplied the materials needed for the experiments. The students would then carry out the experiments in accordance with their design, taking notes on their observations in order to draw conclusions.
6. In relation to the theoretical sections, the students were expected to present to the class the conclusions they drew from their observations and experimental data.
7. After conducting the experiment, the groups attempted to respond to the questions relating to the experiment and the conclusions have been discussed in the classroom.
8. Assessments were carried out following each experiment to determine the extent to which students understood theoretical and scientific information relating to the lessons or experiments carried out.

Furthermore, the selected teacher used to prepare daily lesson plans by using the phases of the 5E lesson plan format for the EG. Engage, explore, explain, elaborate, and evaluate are the phases. For each lesson, all the above steps were structured in to this format. The majority of the guided inquiry based laboratory activities were incorporated in to the exploration phase for the EG. The Control Group (CG) was taught the same topics using the Traditional Laboratory Experiments Enriched Instruction (TLEI) approach, which primarily involved lecture-based teaching and teacher demonstrations. In this method, students acted as passive observers and relied on their textbooks. This approach differed from that used for the Experimental Group (EG) (see Table 2).During the intervention, the researchers used to carry out meetings with the selected teachers at each week ends to discuss on how to prepare daily lesson plan and how to implement the activities of the intervention in the treatment. At the end of the interventions post-test was delivered to all the students in the two groups. Moreover, classroom observation was carried out by the researchers using checklist during the intervention to evaluate whether the instruction was delivered according to the plan or not. The researchers also used to take notes and spent some time to interact with the students, as a participant observer.

**Table 2.**
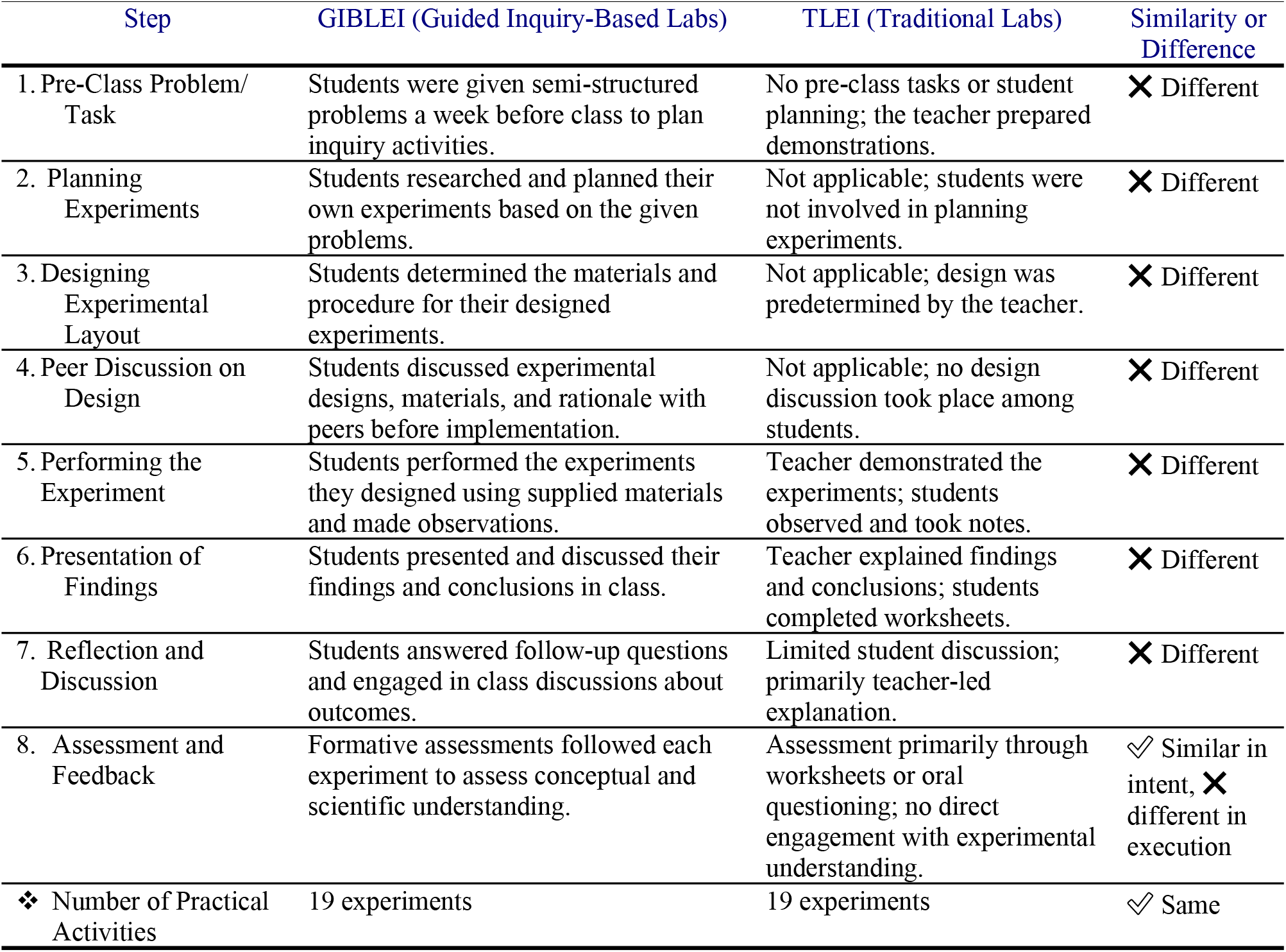
Comparison of GIBLEI and TLEI approaches across the 8-Step Intervention Framework.

### 2.5 Data analysis

Before conducting statistical analyses, the data was assessed to determine whether it satisfied the assumptions for parametric or non-parametric tests. Normality was evaluated using skewness and kurtosis values, where acceptable ranges were set between -2 and +2 (28). Homogeneity of variance was checked using Levene’s test. The results indicated that all variables followed a normal distribution. Tests for homogeneity showed equal variances (p > 0.05) for pretest scores, posttest scores by gender, and the attitude dimension “Biology as a career” (Dimension-4) between the GIBLEI and the TLEI Group (see Table 3). However, significant differences in variances (p < 0.05) were found in the posttest results for overall attitude and other dimensions, including enthusiasm (Dimension-1), biology learning as a course (Dimension-2), and biology as a process (Dimension-3).

**Table 3.** Normality and homogeneity tests for BAQ scores.

| Tests | Independent Variables | Group | Normality test |  | Homogeneity test |  |  |  |
| --- | --- | --- | --- | --- | --- | --- | --- | --- |
|  |  |  | Skewness | Kurtosis | Levene's test |  |  |  |
|  |  |  |  |  | F | df1 | df2 | P |
| **Pretest** | Overall attitude | EG | -0.54 | -0.71 | 3.32 | 1 | 73 | .072 |
|  |  | CG | -0.42 | -0.40 |  |  |  |  |
| **Post-test** | Overall attitude | EG | -0.45 | 0.15 | 21.07 | 1 | 73 | .00 |
|  |  | CG | -0.11 | -0.89 |  |  |  |  |
|  |  | Male(EG) | -0.73 | 0.146 | .110 | 1 | 44 | .742 |
|  |  | Female(EG) | 0.47 | 0.17 |  |  |  |  |
|  |  | Male (CG ) | 0.675 | 0.935 | .128 | 1 | 27 | .723 |
|  |  | Female (CG) | -0.685 | -0.571 |  |  |  |  |
|  | Enthusiasm | EG | -1.242 | 1.179 | 75.79 | 1 | 73 | .00 |
|  |  | CG | -.230 | -1.054 |  |  |  |  |
|  | Biology learning | EG | -.557 | -.335 | 19.30 | 1 | 73 | .00 |
|  |  | CG | -.561 | -.610 |  |  |  |  |
|  | Biology as a process | EG | -.709 | .128 | 24.67 | 1 | 73 | .00 |
|  |  | CG | -.629 | 1.862 |  |  |  |  |
|  | Biology as a career | EG | -.485 | .011 | 1.27 | 1 | 73 | .26 |
|  |  | CG | -.173 | .187 |  |  |  |  |

Based on these findings, appropriate parametric tests were employed. For comparisons involving unequal variances, Welch’s t-test was used, while independent samples t-tests and paired samples t-tests were applied where variances were equal. To measure the magnitude of observed differences, effect sizes were calculated using Cohen’s d (29). Pearson correlation analyses were also conducted to examine the relationships among students’ posttest scores for overall biology attitudes and four subscales (enthusiasm toward biology, biology learning as a course, biology as a process, and biology as a future vocation) separately for the EG and the CG. All analyses were conducted using SPSS version 22.

## 3. Results of the study

### 3.1 Pretest Scores

The pretest results show that there were no statistically significant differences between the EG and CG in their overall attitude toward biology or in any of the four attitude dimensions (enthusiasm, biology learning, biology as a process, and biology as a career), as indicated by the independent samples t-tests (see Table 4). This suggests that both groups had comparable baseline attitudes toward biology prior to the intervention, ensuring the initial equivalence of the groups. Table 5 further explores gender differences within each instructional group. In the EG, no significant gender differences were found across the overall attitude and the four dimensions. However, in the CG, significant gender differences were observed in overall attitude (p = .027), biology learning (p = .021), and biology as a career (p = .005), with female students scoring higher than male students in these areas. These findings indicate that, although the groups were generally equivalent at baseline, some gender-based differences existed within the CG before the intervention.

**Table 4.** Comparison of the overall attitude and its dimensions pretest scores b/n groups (Independent samples t-tests)

| Test | Dependent variables | Group | N | M | SD | T | df | p-value |
| --- | --- | --- | --- | --- | --- | --- | --- | --- |
| **Pretest** | Overall attitude | EG | 46 | 3.96 | .49 | -1.18 | 73 | .241 |
|  |  | CG | 29 | 4.09 | .37 |  |  |  |
|  | Enthusiasm | EG | 46 | 4.07 | .72 | -1.62 | 73 | .108 |
|  |  | CG | 29 | 4.32 | .49 |  |  |  |
|  | Biology learning | EG | 46 | 3.99 | .67 | -.80 | 73 | .426 |
|  |  | CG | 29 | 4.11 | .53 |  |  |  |
|  | Biology as a process | EG | 46 | 3.82 | .61 | -.77 | 73 | .441 |
|  |  | CG | 29 | 3.93 | .61 |  |  |  |
|  | Biology as a career | EG | 46 | 3.99 | .63 | -.37 | 73 | .709 |
|  |  | CG | 29 | 4.04 | .65 |  |  |  |

**Table 5.** Comparison of male and female students’ pretest attitudes toward biology within the EG and CG across the overall and the four attitude dimensions (independent samples t-tests)

| Test | Dependent variables | Group | Gender | N | Mean | SD | t | Df | p-value |
| --- | --- | --- | --- | --- | --- | --- | --- | --- | --- |
| **Pretest** | Overall attitude | **EG** | M | 22 | 3.88 | .51 | -1.14 | 44 | .261 |
|  |  |  | F | 24 | 4.04 | .48 |  |  |  |
|  |  | **CG** | M | 13 | 3.92 | .42 | -2.34 | 27 | .027 |
|  |  |  | F | 16 | 4.23 | .26 |  |  |  |
|  | Enthusiasm | **EG** | M | 22 | 4.05 | .70 | -.14 | 44 | .88 |
|  |  |  | F | 24 | 4.08 | .76 |  |  |  |
|  |  | **CG** | M | 13 | 4.27 | .47 | -.48 | 27 | .633 |
|  |  |  | F | 16 | 4.36 | .53 |  |  |  |
|  | Biology learning | **EG** | M | 22 | 3.85 | .65 | -1.43 | 44 | .158 |
|  |  |  | F | 24 | 4.13 | .67 |  |  |  |
|  |  | **CG** | M | 13 | 3.86 | .55 | -2.45 | 27 | .021 |
|  |  |  | F | 16 | 4.31 | .43 |  |  |  |
|  | Biology as a process | **EG** | M | 22 | 2.70 | 1.92 | -2.36 | 44 | .022 |
|  |  |  | F | 24 | 3.75 | .97 |  |  |  |
|  |  | **CG** | M | 13 | 3.98 | .78 | .346 | 27 | .732 |
|  |  |  | F | 16 | 3.90 | .45 |  |  |  |
|  | Biology as a career | **EG** | M | 22 | 3.90 | .57 | -.91 | 44 | .366 |
|  |  |  | F | 24 | 4.07 | .68 |  |  |  |
|  |  | **CG** | M | 13 | 3.68 | .56 | -3.0 | 27 | .005 |
|  |  |  | F | 16 | 4.34 | .58 |  |  |  |

### 3.2 Overall Attitude Scores

The BAQ post-test mean scores displayed a significant difference between the EG and CG, favoring the EG (t=7.324, p < .05, η2=1.43). This indicates that the GIBLEI approach positively influenced the overall attitude towards biology compared to TLEIs (see Table 6). Consequently, the first null hypothesis (Ho_1_) was rejected.

**Table 6.** BAQ posttest mean scores comparison between the GIBLEI and TLEI groups (Welch’s t-test)

| Group | N | Mean | SD | T | p-value | $\eta^2$ |
| --- | --- | --- | --- | --- | --- | --- |
| EG | 46 | 4.60 | .18 | 7.324 | 0.00 | 1.43 |
| CG | 29 | 4.06 | .36 |  |  |  |

#### 3.3 Attitude Dimensions Analysis

While no significant difference was observed in biology as a career dimension (t=1.81, p >.05, η2=.42), significant differences were found in the post-test mean scores of enthusiasm (t=5.842, p < .05, η2=1.30), biology as a process (t=3.77, p < .05, η2=1.78), and biology learning as a course(t=3.77, p < .05, η2=.82) between the EG and CG, favoring the EG (see Table 7 & 8). This suggests that the GIBLEI approach had a notable impact on students’ perception of enthusiasm, biology learning as a course, and biology as a process compared to traditional methods. Hence, hypothesis two (Ho_2_) was rejected for the former three dimensions, but accepted for the fourth dimension (biology as a career) indicating that the GIBLEI approach did not significantly impact students’ perceptions of biology as a career compared to traditional methods.

**Table 7.** Comparison of attitude dimensions posttest mean scores between groups (Welch’s t-test)

| Test | Variable | Group | N | M | SD | T | p-value | $\eta^2$ |
| --- | --- | --- | --- | --- | --- | --- | --- | --- |
| Post-tests | Enthusiasm | EG | 46 | 4.88 | .14 | 5.842 | 0.00 | 1.30 |
|  |  | CG | 29 | 4.30 | .52 |  |  |  |
|  | Biology learning | EG | 46 | 4.37 | .34 | 3.77 | 0.00 | .82 |
|  |  | CG | 29 | 3.94 | .65 |  |  |  |
|  | Biology as a process | EG | 46 | 4.90 | .09 | 12.58 | 0.00 | 1.78 |
|  |  | CG | 29 | 3.97 | .39 |  |  |  |

**Table 8.** Comparison of attitude dimensions posttest mean scores between groups (independent samples t-tests)

| Test | Variable | Group | N | M | SD | T | p-value | $\eta^2$ |
| --- | --- | --- | --- | --- | --- | --- | --- | --- |
| Post-tests | Biology as a career | EG | 46 | 4.32 | .41 | 1.81 | .074 | .42 |
|  |  | CG | 29 | 4.12 | .52 |  |  |  |

Moreover, Tables 9 and 10 present the Pearson correlations among posttest scores for overall biology attitudes and attitude dimensions in the GIBLEI and TLEI groups, respectively. In both groups, Overall Biology Attitude showed strong positive correlations with Enthusiasm toward biology, Biology learning as a course, and Biology as a future vocation. The correlations were slightly stronger in the EG, particularly between Overall biology attitude and biology learning as a course (r = .806) and Biology as a future vocation (r = .825). Enthusiasm toward biology was positively associated with Biology learning and future vocation in both groups, but not significantly related to Biology as a process. Across both groups, Biology as a process showed weak and mostly non-significant correlations with other variables. These findings suggest that GIBLEI instruction fostered stronger and more interconnected positive attitudes toward biology compared to the traditional TLEI approach.

**Table 9.** Pearson correlations among posttest scores for overall attitudes and attitude dimensions in the EG.

| Variables | 1 | 2 | 3 | 4 | 5 |
| --- | --- | --- | --- | --- | --- |
| 1. Overall Biology Attitude | 1 | .509** | .806** | .333* | .825** |
| 2. Enthusiasm Toward Biology |  | 1 | .333* | .216 | .315* |
| 3. Biology Learning as a Course |  |  | 1 | .160 | .398** |
| 4. Biology as a Process |  |  |  | 1 | .163 |
| 5. Biology as a Future Vocation |  |  |  |  | 1 |

Note: N = 46, $p < .05$ (2-tailed); \*\* $p < .01$ (2-tailed)

**Table 10.** Pearson correlations among posttest scores for overall attitudes and attitude dimensions in the CG.

| Variables | 1 | 2 | 3 | 4 | 5 |
| --- | --- | --- | --- | --- | --- |
| 1. Overall Biology Attitude | 1 | .793** | .812** | .273 | .740** |
| 2. Enthusiasm Toward Biology |  | 1 | .582** | .093 | .483** |
| 3. Biology Learning as a Course |  |  | 1 | -.124 | .385* |
| 4. Biology as a Process |  |  |  | 1 | .140 |
| 5. Biology as a Future Vocation |  |  |  |  | 1 |

Note: N = 29, $p < .05$ (2-tailed); \*\* $p < .01$ (2-tailed)

#### Pretest vs. Post-test Analysis

Significant differences were observed between pretest and posttest mean scores in the EG (t = -8.74, p < .05, η2= 0.19), indicating an improvement in attitude post-intervention (see Table 11). However, there was no significant difference in the CG, implying minimal change with traditional methods. This resulted in the rejection of hypothesis three (Ho_3_) for the EG.

**Table 11.** Comparison of BAQ pre and posttests mean scores in both the EG and CG.

| Group | Test | N | Df | T | p-value | $\eta^2$ |
| --- | --- | --- | --- | --- | --- | --- |
| EG | Pretest | 46 | 45 | -8.74 | 0.00 | 0.19 |
|  | Post-test | 46 |  |  |  |  |
| CG | Pretest | 29 | 28 | .495 | .624 | 0.09 |
|  | Post-test | 29 |  |  |  |  |

#### Gender Analysis

The gender analysis revealed significant differences in student attitudes between male and female students within the CG. At the pretest stage, females outperformed males in overall attitude (M = 4.23 vs. M = 3.92, p = .027), biology learning (M = 4.31 vs. M = 3.86, p = .021), and biology as a career (M = 4.34 vs. M = 3.68, p = .005). These differences persisted into the posttest, where females continued to score higher than males in overall attitude (M = 4.19 vs. M = 3.89, p = .025). In contrast, the EG showed no significant gender differences at pretest (p = .261), indicating a relatively balanced starting point between male and female students (see Table 5 and 12).

**Table 12.** Comparison of male and female students’ post-test attitudes toward biology within the EG and CG across the overall and the four attitude dimensions (independent samples t-tests)

| Test | Dependent variables | Group | Gender | N | Mean | SD | t | Df | p-value |
| --- | --- | --- | --- | --- | --- | --- | --- | --- | --- |
| **Post-test** | Overall attitude | **EG** | M | 22 | 4.58 | .18 | -.78 | 44 | .439 |
|  |  |  | F | 24 | 4.62 | .18 |  |  |  |
|  |  | **CG** | M | 13 | 3.89 | .36 | -2.36 | 27 | .025 |
|  |  |  | F | 16 | 4.19 | .32 |  |  |  |
|  | Enthusiasm | **EG** | M | 22 | 4.83 | .16 | -2.24 | 44 | .030 |
|  |  |  | F | 24 | 4.92 | .11 |  |  |  |
|  |  | **CG** | M | 13 | 4.12 | .56 | -1.71 | 27 | .099 |
|  |  |  | F | 16 | 4.44 | .46 |  |  |  |
|  | Biology learning | **EG** | M | 22 | 4.33 | .29 | -.80 | 44 | .423 |
|  |  |  | F | 24 | 4.41 | .38 |  |  |  |
|  |  | **CG** | M | 13 | 3.67 | .77 | -2.04 | 27 | .052 |
|  |  |  | F | 16 | 4.15 | .46 |  |  |  |
|  | Biology as a process | **EG** | M | 22 | 4.90 | .07 | .21 | 44 | .830 |
|  |  |  | F | 24 | 4.89 | .10 |  |  |  |
|  |  | **CG** | M | 13 | 3.95 | .38 | -.19 | 27 | .847 |
|  |  |  | F | 16 | 3.98 | .40 |  |  |  |
|  | Biology as a career | **EG** | M | 22 | 4.26 | .42 | -.836 | 44 | .407 |
|  |  |  | F | 24 | 4.37 | .40 |  |  |  |
|  |  | **CG** | M | 13 | 3.93 | .40 | -1.821 | 27 | .080 |
|  |  |  | F | 16 | 4.27 | .57 |  |  |  |

Posttest results for the EG showed that male and female students maintained statistically similar attitudes toward biology (p = .439), with females (M = 4.62) slightly ahead of males (M = 4.58). A small but significant difference was noted in enthusiasm, favoring females (M = 4.92 vs. M = 4.83, p = .030). However, no statistically significant gender differences were observed in biology learning (p = .423), biology as a process (p = .830), or biology as a career (p = .407), suggesting balanced posttest outcomes across these dimensions. In the CG, gender differences persisted or approached significance, with females continuing to score higher than males in enthusiasm (M = 4.44 vs. M = 4.12, p = .099), biology learning (M = 4.15 vs. M = 3.67, p = .052), and biology as a career (M = 4.27 vs. M = 3.93, p = .080) (see Table 12).

Analysis of pre-post mean differences showed that in the EG, both genders improved across all attitude dimensions (see Table 13). Males showed an increase of +0.70 in overall attitude and +0.78 in enthusiasm, while females improved by +0.58 and +0.84 respectively. Biology learning as a course means scores also rose for males (+0.48) and females (+0.28). A significant increase was seen in biology as a process for both genders: males improved by +2.20 points and females by +1.14 points. In contrast, the TLEI group exhibited declines in several areas. Males’ overall attitude dropped slightly (−0.03), as did females’ (−0.04). Biology learning as a course scores fell for both males (−0.19) and females (−0.16), and enthusiasm declined among males (−0.15) while females saw only a small gain (+0.08). In biology as a career, males had a minor improvement (+0.25) and females a small decrease (−0.07).

**Table 13.** Pre/Post Comparison within Gender Groups *(Overall Attitude and Each Dimension)*

| Dependent Variable | Group | Gender | Pretest Mean (M) | Posttest Mean (M) | Mean Difference (Post - Pre) |
| --- | --- | --- | --- | --- | --- |
| **Overall Attitude** | EG | Male | 3.88 | 4.58 | +0.70 |
|  | EG | Female | 4.04 | 4.62 | +0.58 |
|  | CG | Male | 3.92 | 3.89 | -0.03 |
|  | CG | Female | 4.23 | 4.19 | -0.04 |
| **Enthusiasm** | EG | Male | 4.05 | 4.83 | +0.78 |
|  | EG | Female | 4.08 | 4.92 | +0.84 |
|  | CG | Male | 4.27 | 4.12 | -0.15 |
|  | CG | Female | 4.36 | 4.44 | +0.08 |
| **Biology Learning** | EG | Male | 3.85 | 4.33 | +0.48 |
|  | EG | Female | 4.13 | 4.41 | +0.28 |
|  | CG | Male | 3.86 | 3.67 | -0.19 |
|  | CG | Female | 4.31 | 4.15 | -0.16 |
| **Biology as a Process** | EG | Male | 2.70 | 4.90 | +2.20 |
|  | EG | Female | 3.75 | 4.89 | +1.14 |
|  | CG | Male | 3.98 | 3.95 | -0.03 |
|  | CG | Female | 3.90 | 3.98 | +0.08 |
| **Biology as a Career** | EG | Male | 3.90 | 4.26 | +0.36 |
|  | EG | Female | 4.07 | 4.37 | +0.30 |
|  | CG | Male | 3.68 | 3.93 | +0.25 |
|  | CG | Female | 4.34 | 4.27 | -0.07 |

## 4. Discussion and Limitations of the findings

### 4.1 Discussion of the findings

This study investigated the effects of the Guided Inquiry-Based Laboratory Experiments Enriched Instructional (GIBLEI) approach on Grade 10 students’ attitudes toward biology, compared to the Traditional Laboratory Experiment Enriched Instructional (TLEI) approach. The findings provided strong evidence favoring the GIBLEI approach. Specifically, analysis using Welch’s t-test revealed that students in the EG (M = 4.60, SD = 0.18) had significantly higher overall attitude scores compared to those in the CG (M = 4.06, SD = 0.36), with a large effect size (ŋ^2^ = 1.43). The rejection of the first null hypothesis (Ho_1_) confirms that inquiry-based instruction is more effective in enhancing students’ general attitudes toward biology than traditional methods. This is consistent with previous studies that highlight the advantages of inquiry-based environments in promoting student engagement, motivation, and interest in science learning (30;10).

Further examination across the four dimensions of attitude revealed that the GIBLEI intervention significantly enhanced students’ attitudes in three of the four measured dimensions. Enthusiasm toward biology showed a strong increase, with the EG scoring a mean of 4.88 compared to 4.30 in the TLEI group (t = 5.842, p < .001, ŋ^2^ = 1.30), indicating a substantial and statistically significant improvement. Likewise, attitudes toward biology learning as a course improved notably, with means of 4.37 for EG and 3.94 for CG (t = 3.77, p < .001, ŋ^2^ = 0.82), suggesting that inquiry-based labs helped students view biology learning more positively. Even more striking was the enhancement in understanding biology as a process, where the EG achieved a mean of 4.90 versus 3.97 for the CG (t = 12.58, p < .001, ŋ^2^ = 1.78), reflecting the strongest effect of the intervention and emphasizing the power of inquiry-based approaches in fostering scientific thinking. These statistically significant improvements led to the rejection of the second null hypothesis (Ho_2_) for these three dimensions. However, when it came to viewing biology as a potential career, no significant difference was observed (M = 4.32 vs. 4.12; t = 1.81, p = .074, ŋ^2^ = 0.42), supporting the acceptance of Ho_2_ in this area. This finding highlights a critical limitation: although GIBLEI can boost immediate academic attitudes, it does not significantly shift students’ longer-term career intentions. This result is consistent with broader research showing that career aspirations are influenced by a complex mix of factors such as mentorship, societal expectations, sustained exposure to role models, and access to real-world STEM experiences (31). Particularly in the Ethiopian context, where sociocultural norms and the limited visibility of female scientists remain significant barriers (32), affecting students’ career trajectories requires more sustained, multi-faceted interventions beyond classroom instruction. Moreover, in the TLEI group, the post-test mean score for students’ attitudes toward biology as a course was lower than their pretest score, indicating a decline in their perception of biology learning after the intervention.

The significant decrease in student attitudes toward biology learning in the CG can be explained by the rigid, non-engaging nature of traditional laboratory activities, which limited opportunities for exploration, critical thinking, and personal relevance. This procedural, passive approach likely made biology seem less interesting and diminished students’ perceived value of the subject over time. Prior research supports this interpretation, showing that teacher-centered methods reduce student enthusiasm and engagement in science learning (33). In Ethiopia, where traditional lecture-based instruction dominates, more than half of secondary school students report that science lessons feel boring and irrelevant, reinforcing biology’s reputation as difficult and uninspiring (32). Importantly, correlation analysis in this study revealed a strong positive relationship between students’ attitudes toward biology learning and their overall attitudes toward biology (r = .812, p < .001). The decline in students’ perceptions of biology learning under the TLEI approach appears to be closely linked to an overall drop in their attitudes toward biology. Since positive attitudes toward science are strongly associated with continued interest and persistence in STEM fields, this trend raises concerns about students potentially losing interest in STEM careers. These findings underscore the urgent need for more interactive, inquiry-based instructional methods like the GIBLEI approach to maintain student engagement and promote long-term participation in STEM.

Paired samples t-tests confirmed the effectiveness of the GIBLEI approach. Students in the EG showed a significant positive shift in attitudes from pretest (M = 3.96, SD = 0.49) to posttest (M = 4.60, SD = 0.18) (t = -8.74, p < .001, ŋ^2^ = 0.19), while the TLEI group showed no significant improvement (t = 0.495, p = .624, ŋ^2^ = 0.09). The positive changes in the EG align with broader research on Inquiry-Based Learning (IBL), which emphasizes how active, student-centered environments enhance motivation and attitudes (8). This is particularly important in resource-constrained settings, where linking abstract biological concepts to real-world experiences helps maintain student engagement (34).

The improvements observed in the EG can be explained by the instructional design that emphasized active inquiry, student collaboration, problem-solving, and critical thinking. Students in the EG were not passive recipients of information but active constructors of knowledge, involved in forming hypotheses, conducting investigations, and analyzing results. This “hands-on, minds-on” approach created an environment where learning was not only meaningful but also personally relevant, leading to enhanced enthusiasm and a more positive perception of biology learning and biological processes(30;35). Consequently, it can be inferred that for secondary school students, instructional strategies that prioritize exploration, autonomy, and meaningful engagement are far more effective in cultivating positive attitudes toward science learning.

The gender analysis provided important insights into how instructional approaches influenced students’ attitudes toward biology. In the EG, posttest results showed minimal gender differences across four attitude dimensions: overall attitude (M = 4.58 vs. 4.62; p = .439), biology learning (M = 4.33 vs. 4.41; p = .423), biology as a process (M = 4.90 vs. 4.89; p = .830), and biology as a career (M = 4.26 vs. 4.37; p = .407). Only the enthusiasm dimension showed a slight but statistically significant difference favoring females (M = 4.92 vs. 4.83; p = .030). These findings align with previous research suggesting that inquiry-based instruction promotes equitable engagement across genders. Erbas & Yenmez (36) found that inquiry-oriented pedagogies support collaborative problem-solving and critical thinking, helping to minimize traditional gender biases common in teacher-centered classrooms. Similarly, Njoku & Nwagbo (37) reported that inquiry-based environments foster active participation among both boys and girls, reducing typical gender gaps in science education. Although GIBLEI minimized gender disparities, the slight female advantage in enthusiasm suggests that some gendered dynamics in STEM attitudes may persist even under inquiry-based instruction. This pattern is consistent with King (38), who observed that female students often show higher affective engagement in interactive, student-centered environments. Gan et al. (39) further explain that females may value the relational and interactive aspects of learning more highly, contributing to greater enthusiasm in such settings.

In contrast, the TLEI group demonstrated persistent and, in some cases, widening gender disparities. At posttest, female students scored significantly higher than males in overall attitude (M = 4.19 vs. 3.89, p = .025) and biology learning (M = 4.15 vs. 3.67, p = .052). Moreover, males in the CG experienced notable declines in enthusiasm (−0.15) and biology learning (−0.19). These findings reflect prior work by Vlckova et al.(40), who emphasized that traditional, lecture-dominated methods often fail to engage male students, leading to disengagement, particularly in disciplines requiring active cognitive engagement like science.

The pre-post mean comparisons reinforce the stark differences in the effectiveness of the two instructional approaches. In the EG, males showed significant improvements in overall attitude (+0.70) and enthusiasm (+0.78), closely mirrored by females (+0.58 and +0.84, respectively). Improvements in understanding biology as a process were particularly striking, with male students improving by +2.20 points and females by +1.14 points. Such gains underscore the capacity of inquiry-based laboratory instruction not only to elevate understanding but to democratize the learning experience across genders (41). By contrast, the TLEI group’s stagnation or regression suggests that traditional instruction may reinforce or exacerbate pre-existing gender disparities. Research by Antonio & Prudente (42) similarly found that rote, textbook-driven learning disproportionately demotivates male students, who often respond better to dynamic, problem-based learning environments.

These contrasting outcomes highlight the critical role of instructional design in shaping students’ attitudes and engagement. The success of the GIBLEI approach aligns with constructivist theories of learning (43), which argue that knowledge is best acquired through active exploration and social interaction rather than passive reception. Inquiry-based learning structures provide opportunities for students to investigate, hypothesize, test, and reflect, engagement processes that naturally appeal across gender lines when properly scaffolded (21). Additionally, equitable learning environments foster not only cognitive growth but also a sense of belonging, which is crucial for underrepresented groups in STEM (44). The GIBLEI model’s ability to create balanced gains across genders suggests that inquiry-based strategies not only promote knowledge acquisition but also help in mitigating systemic attitudinal barriers that can discourage students from pursuing science further.

### 4.2 Limitations

Variations in resource availability and teacher-related variables could have influenced the research outcomes. Conducting the study in natural school environments, where classes were organized by administrators, may also limit the generalizability of the results to other settings. To address these limitations and enhance the validity of future research, several strategies can be considered. These include standardization of resources, enhanced teacher training and monitoring, implementation of randomized controlled trials, longitudinal studies to track long-term effects, and collaborative research partnerships with stakeholders to ensure alignment with educational priorities. By implementing these strategies, researchers can overcome the identified limitations and contribute to advancing knowledge on effective instructional approaches in biology education.

## 5. Conclusions and Implications

### 5.1 Conclusions

This study strongly demonstrates that the Guided Inquiry-Based Laboratory Experiments Enriched Instructional (GIBLEI) approach markedly improves Grade 10 students’ attitudes toward biology compared to the Traditional Laboratory Experiment Enriched Instructional (TLEI) approach. Students exposed to GIBLEI demonstrated higher overall attitude scores, particularly in enthusiasm toward biology, perceptions of biology learning, and understanding biology as a scientific process. These findings confirm that inquiry-based, student-centered instruction is more effective than traditional, teacher-centered methods in fostering positive attitudes toward science learning.

While GIBLEI substantially improved academic attitudes, it had limited impact on students’ career aspirations in biology. This may be due to the curriculum’s lack of explicit discussions about biology-related careers, such as those in botany, soil science, or environmental science. The study’s timing, shortly after the COVID-19 pandemic disruptions, may have further influenced students’ career-related attitudes, given the uncertainty regarding future prospects during that period. Additionally, the GIBLEI approach promoted gender equity, minimizing traditional disparities in science engagement between male and female students, while the TLEI method appeared to reinforce or widen these gaps.

The results highlight the need for Ethiopian secondary schools and similar educational contexts to transition from rigid, procedural laboratory instruction to inquiry-based strategies that emphasize active participation, critical thinking, and real-world relevance. Teachers should be supported through targeted professional development to implement inquiry-based methods effectively. By creating more engaging, equitable learning environments, it is possible to not only improve students’ attitudes toward biology but also to strengthen their long-term interest in STEM fields.

### 5.2 Implications

The findings of this study provide compelling evidence for the efficacy of the Guided Inquiry-Based Laboratory Experiments enriched Instructional (GIBLEI) approach in enhancing secondary school students’ attitudes toward biology. Significant improvements in enthusiasm, perception of biology as a course, and understanding of biology as a process underscore its capacity to foster active engagement and intellectual curiosity. The incorporation of GIBLEI activities into biology curricula at both national and institutional levels can create a more interactive and student-centered learning environment. To optimize implementation, it is essential to equip educators with the necessary skills through targeted teacher training programs that emphasize collaborative and interactive experimental design.

However, the study revealed that the GIBLEI approach has a limited impact on students’ perceptions of biology as a career. Addressing this limitation requires the integration of career-oriented interventions, such as guest lectures by professionals, field visits, and career guidance sessions, to broaden students’ awareness of biology-related career pathways. These enhancements could complement the existing strengths of GIBLEI and ensure that its influence extends beyond academic attitudes to career aspirations. Furthermore, the gender-neutral outcomes of the GIBLEI approach highlight its capacity to foster equity in science education, making it an effective strategy for reducing disparities in STEM participation.

Policymakers and educational stakeholders should prioritize the adoption of GIBLEI by allocating resources for laboratory-based learning, promoting teacher development programs, and ensuring equitable access to inquiry-based methodologies in resource-limited settings. Future research should investigate the long-term effects of the GIBLEI approach on students’ career trajectories and explore methods for seamlessly integrating career-focused elements into inquiry-based pedagogy. By addressing these dimensions, the GIBLEI approach can play a pivotal role in advancing science education and inspiring the next generation of scientists.

## Acknowledgment

We are grateful to the department of Biology, Hawassa University, for unlimited material and official support.

## Author contributions

Every author contributed to the article’s design, data collection, interpretation, writing, and critical revision.

## Data availability

The authors can provide the data produced during this study upon request.

## Declarations

### Ethical Statement

The Research Review Ethics Committee of Hawassa University approved this study under Reference No. CNCS-REC029/22. Additionally, all participants gave their written consent to take part in the research. The study was conducted with written informed consent of the school director, teachers, and students.

